# The topological shape of gene expression across the evolution of flowering plants

**DOI:** 10.1101/2022.09.07.506951

**Authors:** Sourabh Palande, Joshua A.M. Kaste, Miles D. Roberts, Kenia Segura Abá, Carly Claucherty, Jamell Dacon, Rei Doko, Thilani B. Jayakody, Hannah R. Jeffery, Nathan Kelly, Andriana Manousidaki, Hannah M Parks, Emily M Roggenkamp, Ally M Schumacher, Jiaxin Yang, Sarah Percival, Jeremy Pardo, Aman Y Husbands, Arjun Krishnan, Beronda L Montgomery, Elizabeth Munch, Addie M Thompson, Alejandra Rougon-Cardoso, Daniel H Chitwood, Robert VanBuren

## Abstract

Since they emerged ~125 million years ago, flowering plants have evolved to dominate the terrestrial landscape and survive in the most inhospitable environments on earth. At their core, these adaptations have been shaped by changes in numerous, interconnected pathways and genes that collectively give rise to emergent biological phenomena. Linking gene expression to morphological outcomes remains a grand challenge in biology, and new approaches are needed to begin to address this gap. Here, we implemented topological data analysis (TDA) to summarize the high dimensionality and noisiness of gene expression data using lens functions that delineate plant tissue and stress responses. Using this framework, we created a topological representation of the shape of gene expression across plant evolution, development, and environment for the phylogenetically diverse flowering plants. The TDA-based Mapper graphs form a well-defined gradient of tissues from leaves to seeds, or from healthy to stressed samples, depending on the lens function. This suggests there are distinct and conserved expression patterns across angiosperms that delineate different tissue types or responses to biotic and abiotic stresses. Genes that correlate with the tissue lens function are enriched in central processes such as photosynthetic, growth and development, housekeeping, or stress responses. Together, our results highlight the power of TDA for analyzing complex biological data and reveal a core expression backbone that defines plant form and function.

**Significance statement:** A grand challenge in biology is to link gene expression to phenotypes across evolution, development, and the environment, but efforts have been hindered by biological complexity and dataset heterogeneity. Here, we implemented topological data analysis across thousands of gene expression datasets in phylogenetically diverse flowering plants. We created a topological representation of gene expression across plants and observed well-defined gradients of tissues from leaves to seeds, or from healthy to environmentally stressed. Using this framework, we identified a core and deeply conserved expression backbone that defines plant form and function, with key patterns that delineate plant tissues, abiotic, and biotic stresses. Our results highlight the power of topological approaches for analyzing complex biological datasets.

## Introduction

Over 300,000 gene expression datasets have been collected for thousands of diverse plant species spanning over 900 million years of evolution (1). This wealth of publicly available datasets spans ecological niches, species, developmental stages, tissues, stresses, and even single cells, providing a largely untapped reservoir of biological information. These diverse datasets provide an opportunity to link insights from various biological disciplines, including ecology, development, physiology, genetics, evolution, biochemistry, and cell biology through a common computational and mathematical framework. These gene expression datasets have been analyzed individually for specific experiments and hypotheses, but large-scale meta-analyses across the publicly available expression datasets are largely nonexistent for plants.

Beyond a common currency that links the subdisciplines of biology, gene expression links its emergent levels. Below gene expression, the genome gives rise to transcriptional networks and protein interactions that are directly responsible for the complexity of gene expression. Above it, gene expression orchestrates cell-specific expression and the development of the organism itself, impacting phenotypes ranging from physiology to plasticity that propagate further to the population, community, and ecological levels. These features, from molecular (DNA, promoter sequences, -omics datasets) to the organismal, population, and ecological levels (life history traits, climatic data from species distributions, etc.) have been used in the past as labels and predicted outputs of machine learning models(2, 3). The structure—the shape—of gene expression in flowering plants is therefore a constraint that is formed by and impacts biological phenomena below and above it, respectively.

Just as we can look upon the shape of a leaf and derive insights into how it functions from multiple perspectives (developmental, physiological, and evolutionary), we can visualize the shape of any type of data using a Mapper graph (4). The Mapper algorithm takes as input a filter function which describes a biological aspect of the data and uses mathematical ideas of shape to return a graph that reveals the underlying structure of the data. Even abstract data types like gene expression datasets therefore have a shape that we can visualize and derive insights from. For instance, Nicolau et al. (2011) visualized the structure of breast cancer gene expression, identifying two distinct branches with differing underlying genotypes and prognostic outcomes that traditional statistical and bioinformatic approaches fail to resolve (5). This structure was revealed using a pairwise correlation distance matrix as input and modeling of the residuals of each sample from a vector of healthy gene expression as a measure of disease severity. In a second example, using a lens of developmental stage on single cell RNASeq data, Rizvi et al. (2017) visualized the underlying structure of gene expression during murine embryonic stem cell differentiation, revealing transient states as well as asynchronous and continuous transitions between cell types (6). In both examples, Mapper allowed the shape of data, through a selected lens, to be visualized. The resulting topology of the graph—in the form of loops, branch points, or flares—allowed previously hidden structures to be seen and novel insights to be derived.

Gene expression across plants has a shape. This shape guides gene expression along evolutionary, developmental, and environmental trajectories, leading to innovations that have marked the successful adaptation and proliferation of plant species. To visualize this shape is to better understand what transcriptional profiles are possible, and to know the boundaries or constraints that permit or limit gene expression. Here, we analyzed publicly available gene expression profiles across diverse flowering plant families and visualized the underlying structure of gene expression in plants as a graph using the Mapper algorithm. We identified unique topological shapes of plant gene expression when viewed through lenses that delineate different tissue or stress responses. These complex, emergent patterns were largely hidden by biological complexity and sample heterogeneity.

## RESULTS

### A representative catalog of flowering plant gene expression

The vast number of gene expression datasets in plants provides a unique opportunity to search for patterns of conservation and divergence throughout angiosperm evolution, across developmental time, tissues, and stress response axes. Previous studies have tried to find common signatures that define different plant tissues or responses to abiotic/biotic stresses, but these have been limited in species breadth (7), depth (8), or had limited downstream analyses (9). Here, we reanalyzed public expression data on the NCBI sequence read archive (SRA) and applied a topological data analysis method to map the shape of gene expression in plants. We included 54 species that captured the broadest phylogenetic diversity within angiosperms while maximizing the breadth of expression at the tissue and stress level (Figure 1a). This includes 44 eudicots across 13 families and 9 monocot species across 2 families, as well as the basal angiosperm *Amborella trichocarpa*. Raw reads were downloaded, cleaned, and reprocessed through a common RNAseq pipeline to remove artifacts related to the different algorithms and downstream analyses used by each group. After filtering datasets with low read mapping, our final set of expression data includes 2,671 samples across seven distinct developmental tissues and nine stress classifications for 54 species.

**Figure 1.**
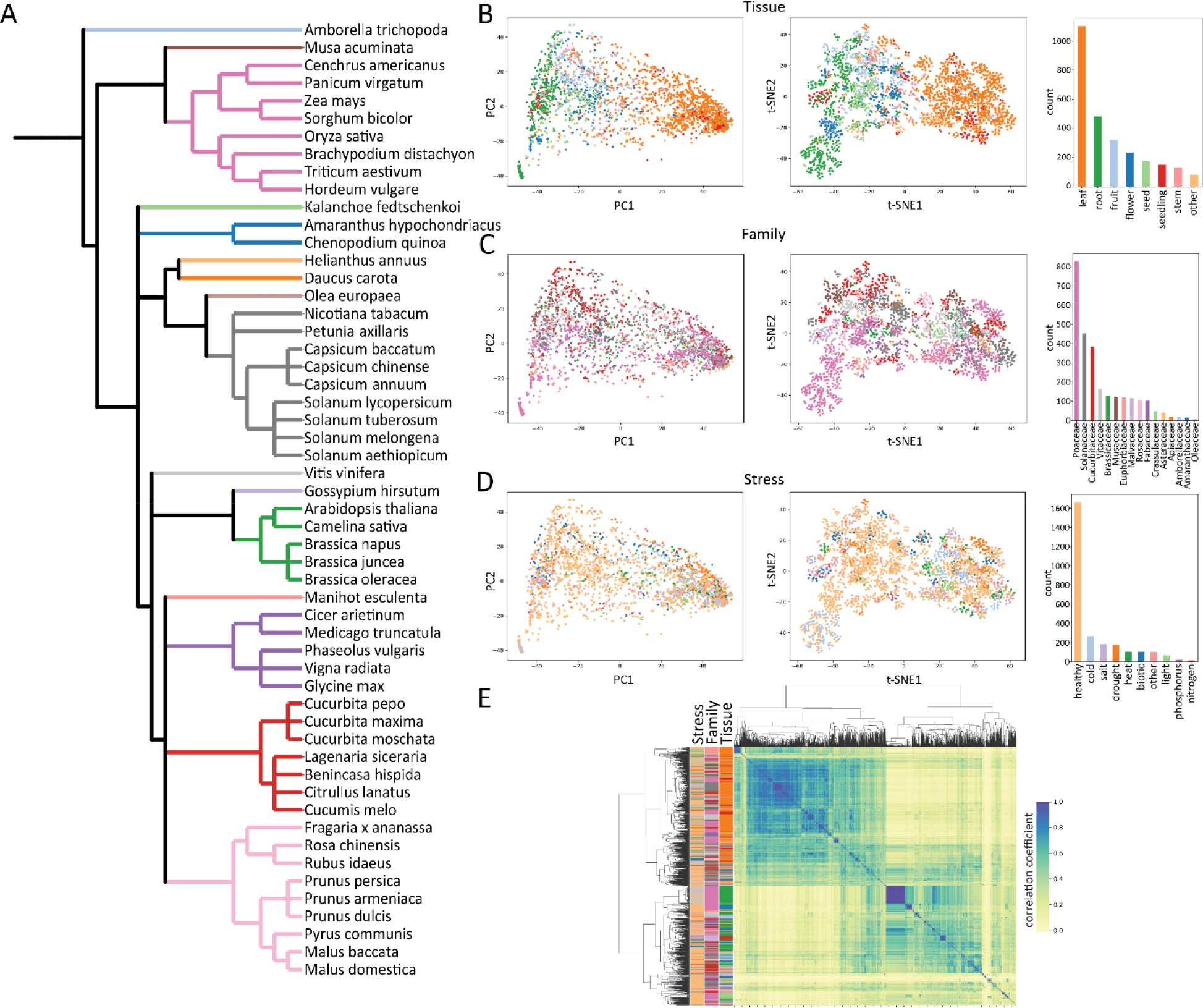
Dimensional space of plant gene expression across evolution, development, and stress. (A) Representative phylogeny of the 54 plant species included in this study. Nodes (species) are colored by plant family as denoted in Figure 1c. Dimensionality reduction of all samples by principal components (left) and t-SNE (right) are shown for tissue type (B), plant family (C), and abiotic/biotic stress (D). Individual samples are quantified and colored by tissue, family, and stress as shown in the respective bar plots. (E) Hierarchical clustering of samples with various biological features highlighted (stress, family, tissue).

To facilitate comparisons of gene expression across species, we limited our analysis to a set of 6,328 orthologous low copy genes that were conserved across all 54 plant species using Orthofinder (10). These sets of orthologous genes or orthogroups are mostly single copy in our diploid species and scale with ploidy in polyploid species. The orthogroups are conserved across a diverse selection of Angiosperm lineages and correspond to well-conserved biological processes. Gene ontology (GO) term enrichment analysis on the Arabidopsis thaliana loci associated with these orthogroups show enrichment for basic biological functions like “DNA replication initiation” and “tRNA methylation” at the top of the list of enriched GO terms, as well as functions specific to photosynthetic organisms like “photosystem II assembly,” and “tetraterpenoid metabolic process.” Although the remaining orthogroups contain significant biological information, they were excluded from analysis as multigene families typically have diverse functions with divergent expression profiles that would conflate downstream comparative analyses.

The transcript per million (TPM) counts were summed for all genes within an orthogroup for a given species, and merged into a single dataframe to create a final matrix of 6,335 orthologs by 2,671 samples. Principal component analysis (PCA) (11) and t-distributed stochastic neighbor embedding (t-SNE) (12) based dimensionality reduction show some separation of samples by different biological factors (Figure 1). The sample space is most clearly delineated by tissue, where both PC1 (explaining 25.4% variation) and t-SNE1 separate the samples into a gradient from root to leaf tissues with other plant tissues sandwiched in between (Figure 1b, 1d). This distribution largely correlates with tissue function, as the sink tissues of flowers, seeds, and fruits resolve closer to the root samples along t-SNE1 and PC1. No tissue type is separated fully by either dimensionality reduction approach. Samples from the 16 plant families are distributed throughout the dimensional space, suggesting family or species level traits are not masking emergent features of distinct tissues (Figure 1c). Interestingly, abiotic and biotic stresses are similarly distributed throughout the dimensional space, with no clear grouping of the same stress across species or individual experiments. This could be due to intrinsic differences in how individual species respond to stress or to differences in the way stress experiments are carried out by different research groups.

### Topological data analysis and the shape of plant gene expression

Traditional dimensionality reduction and hierarchical clustering provided some degree of separation, but they were unable to delineate samples by stress or to identify expression patterns related to biological function. This may be related to residual heterogeneity, noise, or because of the inherent biological complexity that underlies plant evolution and function. To test these possibilities, we used a topological data analysis approach to map the shape of our data. TDA was implemented using Mapper (13), which provides a compact, multi-scale representation of the data that is well suited for visual exploration and analysis. Mapper is particularly well suited for genomics data as these datasets typically have extremely high dimensionality and sparsity (5). To construct mapper graphs from our gene expression data, we created two different lenses of tissue and stress, adopting an approach similar to Nicolau et al (Figure 2a, b). To create the stress lens, we first identified all the healthy samples from the dataset and fit a linear model to them (Figure 2b; see methods). This model serves as the idealized healthy orthogroup expression. We then projected all the samples onto this linear model and obtained the residuals. These residuals measure the deviation of the sample gene expression from the modeled healthy expression and the lens function is simply the length of the residual vector.

**Figure 2.**
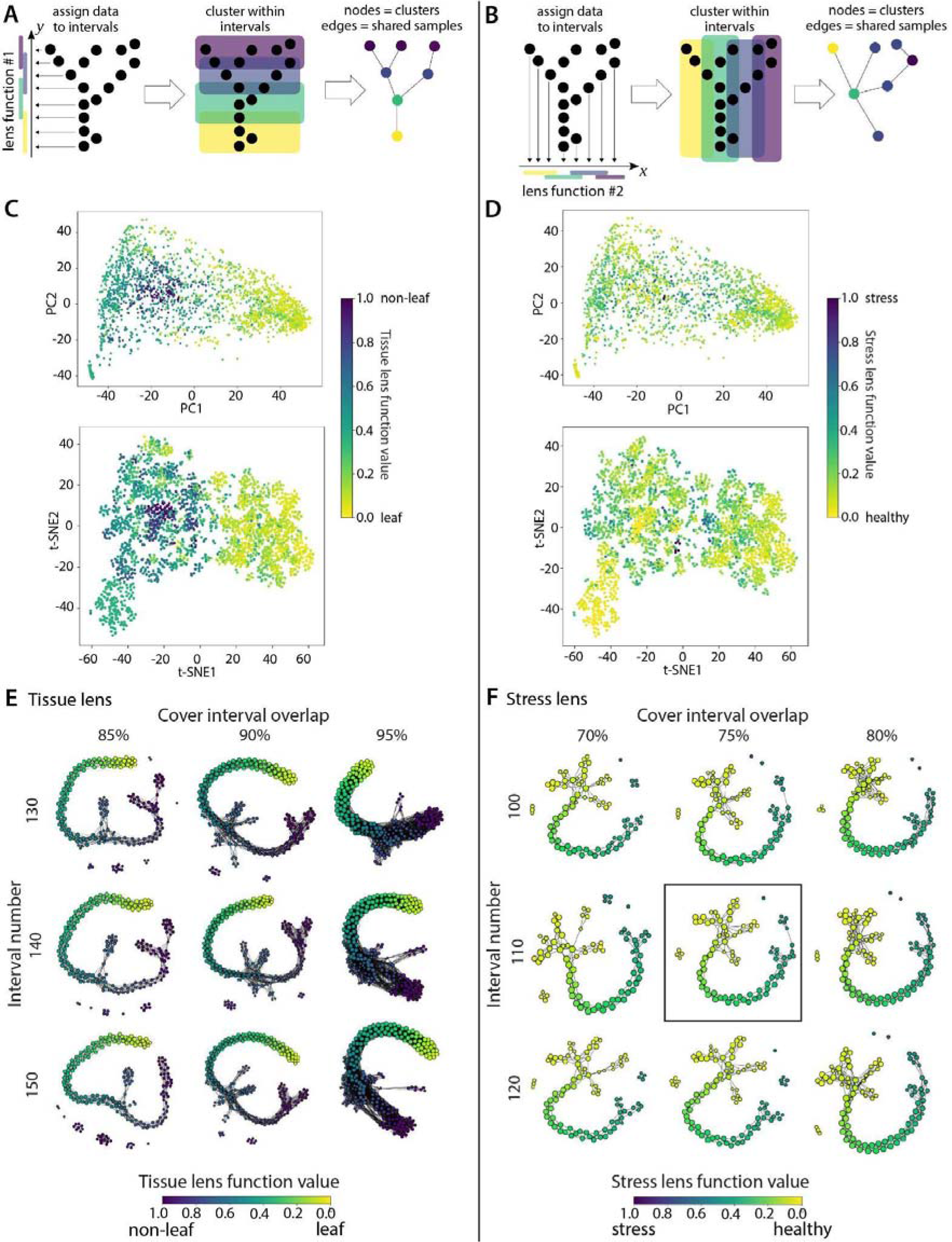
Topology based mapper graphs and the shape of gene expression in plants. Overview of mapper graph construction and lens functions for plant tissue (a) and stress (b). The lens function value of each sample is shown in the principal component (top) and t-SNE (bottom) based dimensional reduction from figure 1 for the tissue (c) and stress lens (d). Mapper graphs across variable cover intervals and interval number for the tissue (e) and stress (f) lens function.

The obvious separation between leaf and root samples in the dimension reduction plots supports a strong photosynthetic vs non-photosynthetic divide. We used this observation to create a binary tissue lens in the same way as the stress lens. We identified all the photosynthetic samples (i.e. leaf tissue), and created an idealized expression profile by fitting a linear model to these expression profiles (Figure 2a). We then projected all the samples onto this linear model and obtained the residuals to establish the lens function by tissue. To define the cover for each lens, we divided the range of the lens function into intervals of uniform length, with the same amount of overlap between adjacent intervals. We experimented with a range of value lengths of the intervals and the size of the overlap to identify the values that produced relatively stable mapper graphs. The clustering was performed using DBSCAN, a commonly used clustering algorithm in Mapper (14).

Overlaying the tissue lens value of each sample over the PCA and t-SNE dimensional space reveals a clear gradient across PC1 and t-SNE1, with the highest lens function values found in seed, fruit, and flower tissues (Figure 2c). For the stress lens function, samples are distributed across the dimensional space, with no obvious correlation between healthy and stressed lens values, similar to the observation from individual abiotic/biotic stresses (Figures 1d and 2d).

Mapper graphs for the tissue and lens functions reflect an emergent and striking topological shape of plant expression (Figure 2e, f). Each node in the Mapper graphs corresponds to a bin of similar RNAseq samples with color representing the average lens value of samples within each node. Edges (connections) show common samples between overlapping bins. Changing the cover interval overlap and interval number has marginal effects on the core graph structure, but changes the shape and connectivity of sparse nodes on the outskirts of the graphs (Figure 2e, f). This central stability highlights the robustness of our input data and significance of the underlying features defining the graph shape (15). The Mapper graphs for both the tissue and stress lens functions show a backbone structure with numerous embedded nodes and flares that form a well-defined gradient from leaf to seed or healthy to stressed respectively. This suggests there are distinct and conserved expression patterns across angiosperms that delineate different tissues or responses to biotic and abiotic stresses.

### Topological shape reflects the underlying biological features of gene expression

To identify and characterize these conserved biological patterns, we first simplified the Mapper graphs into 18 and 17 nodes for the tissue and stress lens function respectively (Figures 3 and 4). The core tissue-based Mapper graph has discrete nodes for each surveyed plant tissue with a gradual transition of leaves (node 1), to roots (2), fruits (11 and 13) and finally seeds (14,15, and 16; Figure 3a). At the 4th node, the Mapper graph proliferates into terminal branches of flower (node 9), stem (10), fruit (12), and mixtures of uncategorized tissue types (5 and 8). RNAseq samples from the 16 angiosperm families are largely dispersed across nodes by tissue, with some notable exceptions (Figure 3b). Most fruit samples are found along the gradient of the core graph structure, but fruits from the rose (Rosaceae) family form a separate node (node 12). Flowers from the eudicot species are mixed with fruit tissues in nodes along the core graph structure, but monocot flowers from the grass family (Poaceae) are found in discrete, branching nodes (9 and 17). The biotic and abiotic stress RNAseq samples are dispersed by tissue across the Mapper graph (Figure 3c), supporting the complexity and heterogeneity of these samples.

**Figure 3.**
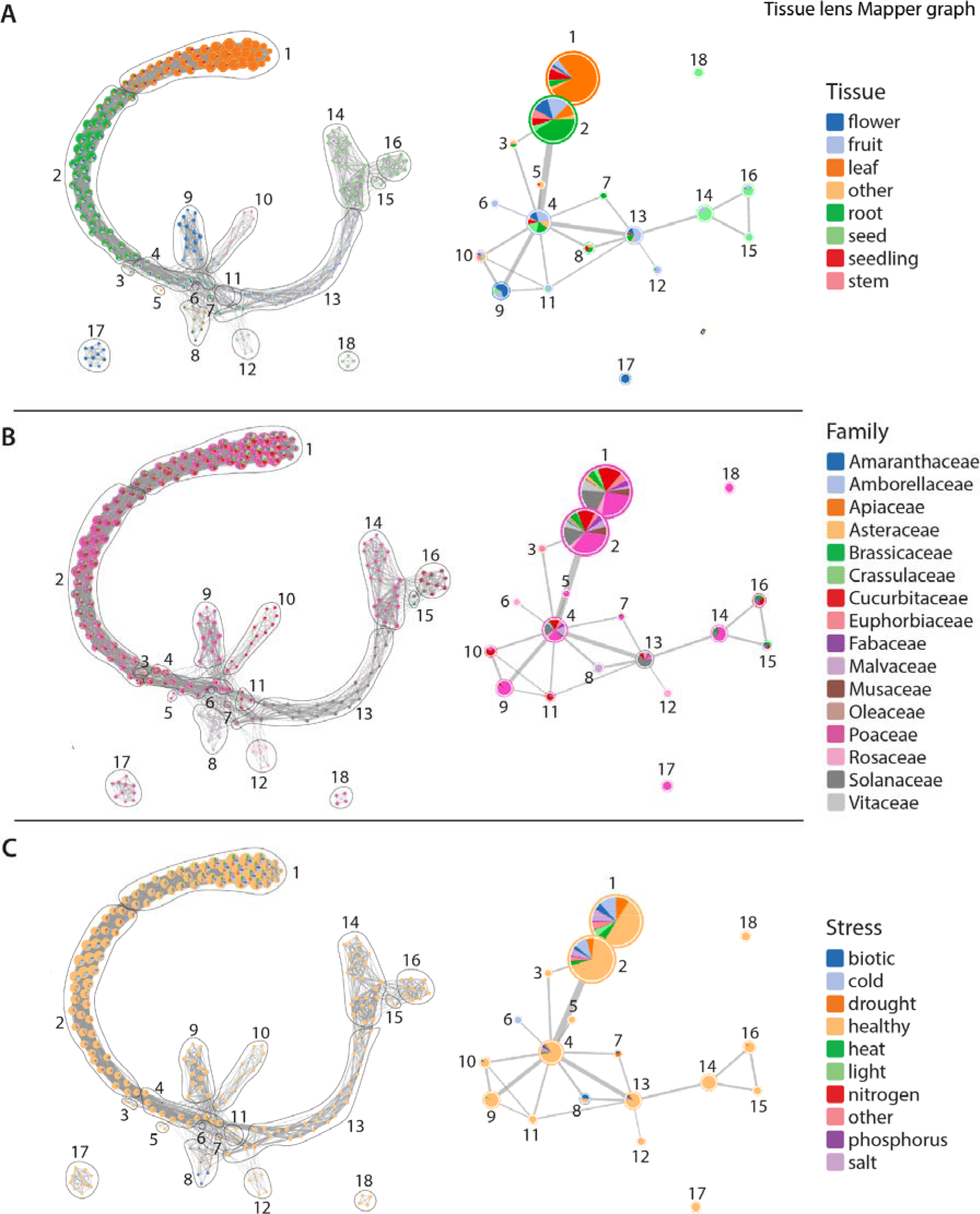
Simplified Mapper graphs detailing the distribution of samples along the stress lens. Nodes along the full Mapper graphs (left) are clustered into simplified Mapper graphs (right) and samples are colored by tissue (A), Family (B), and Stress category (C).

**Figure 4.**
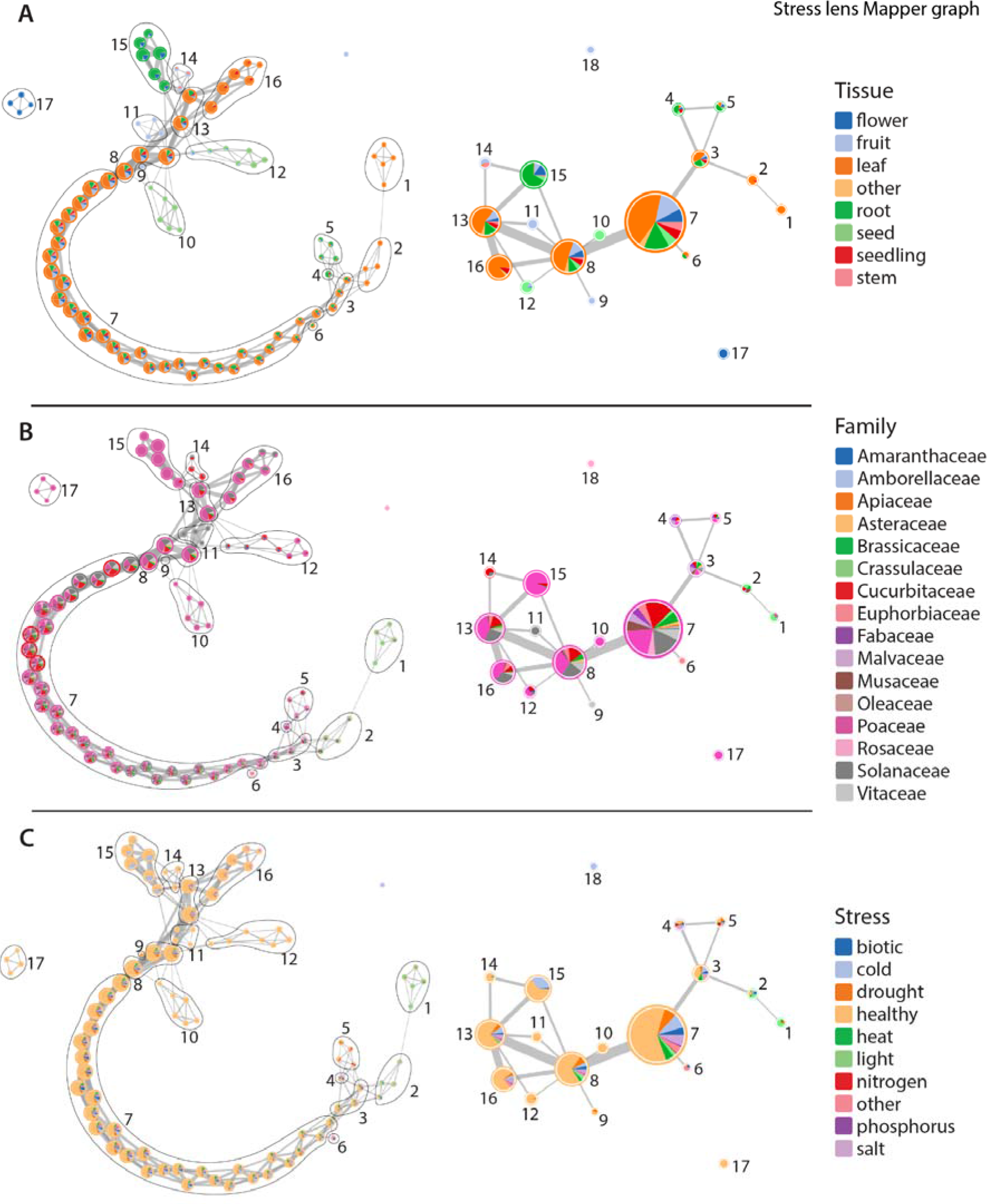
Simplified Mapper graphs detailing the distribution of samples along the stress lens. Nodes along the full Mapper graphs (left) are clustered into simplified Mapper graphs (right) and samples are colored by tissue (A), Family (B), and Stress category (C).

Mapper graphs clearly distinguish tissues across plant taxa but what are the biological features that underlie this topology? We surveyed the expression patterns of the 6,328 orthogroups used to generate our Mapper graphs to see if they are enriched in certain biological processes related to evolutionarily conserved, tissue specific functions. We classified genes as positively or negatively correlated with the tissue lens and conducted gene ontology (GO) enrichment in these groups of genes. We expect negatively correlated genes to be characteristic of leaf gene expression and positively correlated genes to be characteristic of non-leaf gene expression. Supporting this, Mapper graphs and GO terms associated with the tissue-lens correlated genes point to photosynthetic vs. non-photosynthetic metabolism as a key factor in the overall gene expression patterns of plant tissues (Figure 3; Supplemental Dataset 1). Enriched negatively correlated GO terms are mostly related to photosynthesis and include response to red and blue light, chloroplast and thylakoid organization, carotenoid metabolic process, and regulation of photosynthesis among others (Supplemental Dataset 1). Plants and green algae are characterized by a set of well-conserved genes that are not found in non-photosynthetic organisms termed the GreenCut2 inventory (16). Most of the GreenCut2 genes (421 out of 677) are found within the 6,328 orthogroups in our analysis and we tested if these are enriched among correlated genes. Genes from the GreenCut2 inventory are overrepresented in this set of genes, with 26.7% of the tissue-correlated (positively or negatively) genes being in the GreenCut2 resource versus 6.7% of the entire set of orthogroups (Supplemental Table 1). This overrepresentation is even more stark if we delimit our analysis to only the genes negatively correlated with the tissue lens, of which 50.3% are in the GreenCut2 inventory. The overlapping loci between the two sets contain genes encoding protein products involved in various aspects of photosynthesis, including pigment biosynthesis and binding (e.g. AT4G10340, AT1G04620, AT1G44446) (17–19), the operation of the photosynthetic light reactions (e.g. AT4G05180, AT5G44650, AT3G17930) (20–22), or the operation of the Calvin-Benson Cycle (AT1G32060) (23).

Enriched GO terms that are positively correlated with the tissue lens are largely related to housekeeping and core metabolic processes including ubiquitination, macromolecule catabolism, the electron transport chain, peptide biosynthesis and Golgi vesicle-mediated transport among many others (Supplemental Dataset 2). Enriched genes include proteins involved in the TCA cycle, and respiration (e.g., AT1G47420, AT2G18450, AT4G26910) (24–26) and in the development of specific non-photosynthetic tissue types like seeds (e.g. AT2G40170, AT2G38560) (27, 28) and pollen/pollen tubes (e.g., AT2G03120, AT2G41630) (29, 30). However, many of the tissue-lens correlated genes do not intuitively relate to the photosynthetic vs. non-photosynthetic tissue distinction and further examination of these loci on a gene-by-gene basis may shed light on conserved differences between plant tissues.

The simplified Mapper graph from the stress lens has 17 nodes that form a continuous gradation of healthy to stressed tissues (Figure 4). Individual tissue types, regardless of stress condition, are enriched in certain nodes, but are less defined than under the tissue lens (Figure 4a). RNAseq samples related to light and heat stress are found in discrete nodes (1 and 2 respectively) at the terminus of the Mapper graph across all species where this data was available (Figure 4c). Other stress RNAseq samples are found in nodes with healthy tissues but are generally concentrated toward the stress end of the Mapper graph. An interesting exception is a group of cold stressed root samples from the grass (Poaceae) family (node 15). Clustering of distinct stresses within the same node suggests a core stress response conserved across Angiosperms for all abiotic and biotic factors. The gradient of sample distribution from healthy to stressed across the Mapper graph may be related to the severity of stress experienced by plants in each individual experiment.

To explore what constitutes these conserved stress related expression patterns, we searched for GO enrichment of genes that are positively correlated with the stress lens. This group of genes is heavily enriched in functions related to stress including responses to water deprivation, chitin, reactive oxygen species, fungi, wounding, bacteria, and general defense mechanisms (Supplemental Dataset 3). Genes positively correlated with the stress-lens include loci related to the biosynthesis of compounds with diverse stress-related activities like jasmonic acid and jasmonic acid derivatives (AT2G35690, AT2G46370) (31, 32) and ascorbic acid (AT3G09940) (33). Negatively correlated genes are enriched in functions related to growth and reproduction such as DNA replication, mitosis, and rRNA processing, among others (Supplemental Dataset 4). This includes genes involved in regulation of the cell cycle (AT3G54650, AT4G12620, AT2G01120) (34–36), chromatin organization (AT1G15660, AT1G65470) (37, 38), and the development of reproductive structures (AT1G34350, AT2G41670, AT4G27640, AT3G52940) (39–42). This pattern points towards an intuitive distinction between the stressed and unstressed samples in our dataset in terms of their investment in cell proliferation and reproduction. Most of these genes are involved in core biological functions with conserved roles across eukaryotes, and their coordinated perturbation could be predictive of stress responses in diverse lineages.

## DISCUSSION

Genome scale datasets have high dimensionality and even the simplest pairwise experiment has hundreds or thousands of complex and interconnected cellular pathways in dynamic flux between conditions. Comparisons across plant lineages are similarly complex, as each species has its own evolutionary history with thousands of duplicated, lost, or new genes enabling its unique and elegant biology. This complexity presents major challenges for characterizing underlying biological mechanisms and identifying shared and distinct properties across evolutionary timescales. Here, we leveraged the wealth of public gene expression datasets across diverse flowering plants, and used a set of deeply conserved genes to search for patterns of conservation across tissue types, stress responses, and evolution. We first tested traditional dimensionality reduction and clustering-based approaches but found they were largely ineffective and unable to clearly resolve samples. Instead, we used a novel topological framework to compare samples and test for evolutionary conservation.

Topological data analysis has been applied to complex, high dimensionality biological datasets including gene expression profiles correlated with human cancers and other diseases (5, 43, 44). To our knowledge, TDA has not been used for plant science datasets outside of shape (45–47). Flowering plants have tremendous phylogenetic, developmental, phenotypic, and genomic scale diversity, creating additional layers of complexity compared to other lineages. Despite this, Mapper was able to capture hidden and emergent signatures of gene expression at the tissue and stress scales that were missed using traditional approaches. Most developmental tissues or stress responses are not perfectly separated, but instead fall within a gradient along a central shape. The central shape of the tissue lens Mapper graph represents the life cycle of a plant with transitions from the vegetative tissues of leaves and roots to reproductive flowers, fruit, and eventually seeds. Nodes along the Mapper graphs that contain mixtures of tissues such as fruits and flowers, leaves and stems, or even leaves and roots reflect developmental plasticity, heterogeneity, and overlapping functions between different organs. Flowers give rise to fruits and the complex processes of fertilization, seed, and fruit development blur the lines between distinct tissue types. This complexity and interconnectivity is central to biological processes, but is masked by traditional dimensionality reduction approaches which can oversimplify nonlinear datasets.

The stressed and healthy samples are less clearly delineated in the Mapper graphs than samples from different plant tissues. This may reflect artifacts stemming from variation in the severity, duration, or method of applying stresses across different experiments and species. For instance, mildly stressed samples might have expression signatures that mirror healthy tissues with comparatively few differentially expressed genes. Despite this issue, we observed a strong gradient of sample distribution from healthy to stressed across the graph. Distinct stresses were generally found within the same nodes, and genes that were positively correlated with the stress lens show enrichment in classical stress pathways. This includes the core stress responsive hormones jasmonic acid and abscisic acid and their corresponding transcriptional network as well as broader shifts in metabolic processes geared toward defense. Taken together, this suggests that plants have deeply conserved expression signatures across evolution and for different stresses. Abiotic and biotic stress responses are historically viewed as evolutionarily independent and transcriptionally unique, but the topological shape of gene expression points to a shared set of pathways that define if a tissue is healthy or stressed.

Although we observed a deeply conserved pattern of gene expression underlying plant form and function, our analyses capture a snapshot of the evolutionary innovations found in flowering plants. We used a set of conserved genes to enable comparisons of expression across species, and we had to exclude thousands of new or rapidly evolving gene families that are associated with important plant traits. Here, we provide a proof of concept for studying complex biological traits using TDA, and a similar analytical framework could be applied to numerous areas of plant science research and beyond. Compared to the ~300,000 published plant gene expression datasets (1), our study has a somewhat sparse sampling of species and a subset of expressed genes, yet were able to detect a number of hidden trends. TDA of high-resolution sampling over narrower phenotypic spaces such as drought responses in a single species, or tissue divergence across 900 million years of plant evolution could yield transformative insights that were previously overlooked.

## METHODS

### Assembling a representative catalog of flowering plant expression data

We selected species that captured the broadest phylogenetic diversity within angiosperms and species that had a breadth of expression at the tissue and stress level. We also selected only species with a high-quality reference genome, to enable accurate read mapping and downstream comparative genomics. Metadata including species, accession, tissue type, experimental treatments, replicate number, and sequencing platform was collected manually for each sample using the NCBI BioProject and sequence read archives (SRA), as well as the primary data publications (Supplemental Dataset6). Raw RNAseq reads were downloaded from the NCBI SRA and quantified using a pipeline developed in the VanBuren lab to trim, quantify, and identify differentially expressed genes (https://github.com/pardojer23/RNAseqV2). Using a common analytical pipeline helped reduce noise between experiments that used different algorithms in the original publications. Raw Illumina reads from various platforms were first quality trimmed using fastp (v0.23) (48) with default parameters. The quality filtered reads were pseudo-aligned to the corresponding transcripts (gene models) for each species using Salmon (v1.6.0) (49) with the quasi-mapping mode. Transcript level estimates were converted to gene level transcript per million counts using the R package tximport (50).

### Comparing expression across species

To facilitate detailed cross-species comparisons, we first clustered proteins from all 54 species into orthogroups using Orthofinder (v2.3.8) (10). Genomes and proteomes were downloaded for each species from Phytozome v13 (51). Orthofinder was run using default parameters and the reciprocal DIAMOND search (v2.0.11) (52) was used for sequence alignment, and groups of similar proteins were clustered using the Markov Cluster Algorithm. In total, 2,317,289 genes (94% of input genes) were clustered into 86,185 orthogroups across the 54 species. Of these, 33,585 orthogroups are found in only a single species and 7,742 are found in at least 52 out of 54 species. This set of broadly conserved orthogroups was further refined by filtering out orthogroups with an average of > 2 genes per ortholog for the diploid species to avoid including multigene families with diverse functions in the analysis. This set of 6,335 orthogroups was used as a common framework to allow comparison of expression across species. For orthogroups where a species had more than one gene, the total TPM for all genes in that orthogroup was summed and the raw TPM was used for single copy genes. Expression data for each sample across all species were combined into a single expression matrix (Supplemental dataset 7).

### Mathematical basis of topological data analysis

The flexibility of Mapper allows us to apply it to various types of data. Here we will describe the Mapper construction in the simplest setting of point cloud data and then explain how it was applied to the gene expression data.

Consider a point cloud ***X*** ⊂ **R**^d^ equipped with a function *f* : ***X*** → **R**. An open cover of ***X*** is a collection ***U*** = {***U***_i_}_i∈**I**_ of open sets in **R**^d^, such that ***X*** ⊂ ⋃ _i∈**I**_ ***U*_i_**, where **I** is an index set. The 1-dimensional nerve of the cover ***U***, denoted as ***M*** ≔ ***N***_1_(***U***), is called the Mapper graph of (***X***, *f*). In this graph, each open set ***U***_i_ is represented as a vertex ***i***, and two vertices, ***i*** and ***j***, are connected by an edge if and only if the intersection of ***U***_i_ and **U**_j_ is nonempty.

To construct a Mapper graph, we start by defining a cover ***V*** = {***V***_j_}_j∈**J**_ of the image *f*(***X***) ⊂ **R** of *f*, where **J** is a finite index set, by splitting the range of *f*(***X***) into a collection of overlapping intervals. Next, for each ***V***_j_, we identify the subset of points ***X***_j_ in ***X*** such that *f*(***X***_j_) ⊂ ***V***_j_ and apply a clustering algorithm to identify clusters of points in ***X***_j_. The cover ***U*** of ***X*** is the collection of such clusters induced by *f*^−1^(***V***_j_) for each j. Once we have the cover ***U***, we compute its 1-dimensional nerve ***M*** and visualize it in the form of a weighted graph.

For example, consider Figure 2a. The point cloud ***X*** in this case consists of points in the 2-dimensional plane, in the shape of a “Y”. The function *f* simply maps each point to its *y*-coordinate. We divide the range of *f* into 4 overlapping intervals, represented by the four colored segments along the *y*-axis in Figure 2a. For each interval ***V***_j_, the colored rectangles in the center panel of the figure show the subsets of points ***X***_j_ ∈ ***X*** such that ***X***_j_ = *f*^−1^(***V***_j_). Then, we apply clustering to each ***X***_j_ separately to obtain the cover ***U*** of ***X***. The 1-dimensional nerve of ***U***, *i.e*. the mapper graph ***M***, is shown in the rightmost panel. The color of each vertex corresponds to the cover interval it belongs to. Figure 2a illustrates mapper graph construction from the same set of points, but with *x*-coordinate used as the lens. We can observe that the two lens functions produce two slightly different mapper graphs.

### Constructing Mapper graphs and lens functions

To construct mapper graphs from our gene expression data, we create two different lenses, adopting an approach similar to the one used in Nicolau et al. We refer to these lenses as the tissue lens and the stress lens, respectively. To create the stress lens, we first identified all the healthy samples from the dataset and fit a linear model to them. This model serves as the idealized healthy orthogroup expression. Then, we project all the samples (healthy as well as stressed) onto this linear model and obtain the residuals. These residuals measure the deviation of the sample gene expression from the modeled healthy expression. The lens function is simply the length of the residual vector. To define the cover, we divide the range of the lens function into intervals of uniform length, with the same amount of overlap between adjacent intervals. We experimented with a range of values length of the intervals and the size of the overlap to identify the values that produced relatively stable Mapper graphs. The clustering was performed using DBSCAN, a commonly used clustering algorithm for Mapper.

The construction of Mapper graph relies on several user defined parameters: the lens function *f*, the cover ***V***, and the clustering algorithm. Optimizing these parameters is an interesting open problem in TDA research (53). The function *f* plays the role of a lens, through which we look at the data, and different lenses provide different insights (4). The choice of *f* is typically driven by the domain knowledge and the data under consideration. In this study, the data under consideration is very similar to the dataset studied by Nicolau et al. (5). Therefore, we followed similar methods to define the lenses. Our choice of lenses is further justified by the observations from the dimension reduction plots.

The cover ***V*** = {***V***_j_}_j∈**J**_ of *f*(***X***) consists of a finite number of open intervals as cover elements. To define ***V***, we use the simple strategy of defining intervals of uniform length and overlap. Adjusting the interval length and the overlap increases or decreases the amount of aggregation provided by the Mapper graph. The optimal choice was made by visually inspecting Mapper graphs over a range of parameter values. The parameters resulting in the most stable structure were selected. Any clustering algorithm can be employed to obtain the cover ***U***. We use the density-based clustering algorithm, DBSCAN (54), which is commonly used in Mapper because it does not require a priori knowledge of the number of clusters. Instead, DBSCAN requires two input parameters: the number of samples in a neighborhood for a point to be considered as a core point, and the maximum distance between two samples for one to be considered in the neighborhood of the other.

### Functional annotation of orthogroups

The correlation between expression values and tissue- and stress-lens values was calculated for each orthogroup. The top 2.5% most positively and negatively correlated orthogroups for each lens were selected to represent the tissue- or stress-lens correlated orthogroups. Arabidopsis gene IDs were used to identify the overlap between the GreenCut2 (16) inventory with Arabidopsis orthologs in our overall set of orthogroups, as well as our sets of tissue- and stress-lens correlated orthogroups. The *binom_test()* function from SciPy (55) was used to apply one-sided binomial tests to check for enrichment of GreenCut2 loci in the overall, tissue-, and stress-lens correlated orthogroup sets. GO Term enrichment of the sets of genes mapped to orthogroups and correlated with the tissue- or stress-lens was done using GOATOOLS (56). Data on gene function and biochemical reactions associated with specific loci were derived from TAIR (57), KEGG (58), and a genome-scale metabolic model of Arabidopsis metabolism from (59).

## Data availability

The code, metadata, and raw datasets from this project are available on a dedicated GitHub page: https://github.com/PlantsAndPython/plant-evo-mapper.git

## Acknowledgements

This work was funded primarily by an NSF-NRT training grant (NSF 1828149) which established the Integrated training Model in Plant And Compu-Tational Sciences (IMPACTS) program at Michigan State University. This grant funded fellows within this program as well as the project-based curriculum for the Plants and Python Course that formed the backbone of this manuscript.

