## Supplemental Figures/Tables for "The topological shape of gene expression across the evolution of flowering plants"

Supplemental materials

Table S1. Enrichment of GreenCut2 genes in orthogroup-mapped *Arabidopsis thaliana* genes and stress-/tissue- correlated orthogroup-mapped genes. The proportion of GreenCut2 genes in the all the orthogroups used in this study was compared against the proportion of GreenCut2 genes in a list of all *A. thaliana* genes using a one-sided binomial test. The proportion of tissue-lens and stress-lens correlated orthogroup-mapped genes in GreenCut2 was compared against the proportion of GreenCut2 genes in the entire set of orthogroup-mapped genes using one-sided binomial tests. Tissue-correlated genes were hypothesized to be more likely to be in GreenCut2 than a random selection of orthogroup-mapped genes, and the stress-correlated genes were hypothesized to be less likely.

| **Dataset** | **# of Genes in Dataset** | **# of Genes in GreenCut2** | **% GreenCut2** | **p-value** |
| --- | --- | --- | --- | --- |
| **All Arabidopsis Genes** | **27662** | **677** | **2.45** |  |
| **All Orthogroup-Mapped Genes** | **6328** | **421** | **6.65** | **2.76 * 10^-96^** |
| **All Tissue-lens Correlated Genes** | **318** | **85** | **26.7** | **9.18 * 10^-29^** |
| **Stress-lens Correlated Genes** | **318** | **7** | **2.20** | **0.000252** |

Dataset S1 (separate file). GO Term enrichment results on genes negatively correlated with the tissue-lens.

Dataset S2 (separate file). GO Term enrichment results on genes positively correlated with the tissue-lens

Dataset S3 (separate file). GO Term enrichment results on genes positively correlated with the stress-lens

Dataset S4 (separate file). GO Term enrichment results on genes positively correlated with the stress-lens

Dataset S5 (separate file). Overlap between orthogroup-mapped genes and tissue- and stress-lens correlated genes with the GreenCut2 resource (Karpowicz

Dataset S6 (separate file). Metadata of the raw data used in this experiment.

Dataset S7 (separate file). Expression matrix of TPMs for the normalized orthogroups
